# HIV proviral genetic diversity, compartmentalization and inferred dynamics in lung and blood during long-term suppressive antiretroviral therapy

**DOI:** 10.1101/2022.05.26.493568

**Authors:** Aniqa Shahid, Bradley R. Jones, Julia Yang, Winnie Dong, Tawimas Shaipanich, Kathryn Donohoe, Chanson J. Brumme, Jeffrey B. Joy, Janice M. Leung, Zabrina L. Brumme

## Abstract

The lung is an understudied site of HIV persistence. We isolated 882 subgenomic proviral sequences by single-genome approaches from blood and lung from nine individuals on long-term suppressive antiretroviral therapy (ART), and characterized genetic diversity and compartmentalization using formal tests. Consistent with clonal expansion as a driver of HIV persistence, identical sequences comprised between 9% to 86% of within-host datasets, though their location (blood vs. lung) followed no consistent pattern. The majority (77%) of participants harbored at least one sequence shared across blood and lung, supporting the migration of clonally-expanded cells between sites. No participant exhibited genetic compartmentalization so obvious that it was visually apparent in a phylogeny. When formal tests were applied however, two (22%) participants showed modest yet significant support for compartmentalization when analysis was restricted to distinct proviruses per site. This increased to four participants (44%) when considering all within-host sequences. Thus, while a minority of individuals harbor somewhat distinctive proviral populations in blood and lung, these can simply be due to unequal distributions of clonally-expanded sequences. Importantly, the extent of lung proviral diversity on ART strongly reflected that in blood (Spearman ρ = 0.98, p < 0.0001), confirming blood as a strong indicator of total body proviral diversity. Analysis of on-ART proviral diversity in context of pre-therapy viral diversity in two participants revealed marked differences in proviral longevity and dynamics. Whereas one participant’s proviral pool was rich in ancestral sequences that recapitulated more of HIV’s within-host evolutionary history, the other’s largely comprised more contemporary sequences, including ones that re-seeded the reservoir during a three-year treatment interruption. Results highlight the genetic complexity of proviruses persisting in lung and blood during ART, and the uniqueness of each individual’s proviral composition. Individualized HIV remission and cure strategies may be needed to overcome these challenges.

**Author Summary:** HIV persists in the body despite long-term suppressive antiretroviral therapy. Much of our knowledge about the HIV reservoir comes from studying proviruses in blood, but a fundamental question is whether these are distinct from those in tissues. The lung could theoretically engender genetically distinctive HIV populations, but this remains understudied. Our analysis of nearly 900 proviral sequences from blood and lung of nine individuals with HIV receiving long-term therapy revealed substantial within-host heterogeneity, yet some common patterns. Identical sequences (consistent with clonal expansion of infected cells) were observed in everyone, though at different (9-86%) frequencies. Recovery of shared sequences across blood and lung was common (77% of participants). Fewer than half of participants exhibited blood-lung genetic compartmentalization, and only modestly so. Moreover, when present, compartmentalization was often attributable to differential distribution of identical sequences, not the presence of genetically distinctive populations, across sites. Results also revealed marked inter-individual differences in proviral dynamics, as evidenced by the proportion of persisting proviruses that represented ancestral versus more recently-circulating viral sequences. Critically, the extent of within-host proviral diversity in blood correlated strongly with that in lung, indicating that despite inter-individual heterogeneity, blood is a strong indicator of proviral diversity elsewhere in the body.

## Introduction

HIV replicates rapidly during untreated infection [1, 2], during which time the virus disseminates throughout the body, including into multiple tissues [3]. Representatives of these replicating viral populations are continually archived into the reservoir, where they can subsequently persist, even during long-term suppressive antiretroviral therapy (ART) [4]. Much of what we know about the HIV reservoir comes from studying proviruses that persist in blood during long-term ART [5, 6]. In general, the diversity of these proviruses tends to reflect the extent of within-host HIV evolution during untreated HIV infection [7–9], though proviruses archived near the time of ART initiation are often overrepresented [10–12]. Some individuals’ reservoirs are also dominated by clonally-expanded populations of infected cells [13–15]. HIV persistence in tissues however remains less well understood. While some studies have revealed similar proviral landscapes in blood and tissue [16–19], others support the presence of distinct populations [3, 20].

One understudied site of HIV persistence is the lung, even though various lines of evidence identify it as a potentially distinct anatomical HIV reservoir [21, 22]. HIV can persist within alveolar macrophages [23, 24], which are abundant in the lung [25]. These distinctive cells are long-lived [26] and relatively resistant to both apoptosis [27–29] and cytotoxic CD8^+^ T-cells [30], and a number of studies have demonstrated a higher proviral burden in pulmonary compared to blood cells [23, 31]. Unique tissue environments can facilitate the evolution of viral populations that differ from one another genetically. This could happen if founder virus(es) entering into such environments subsequently evolve ‘locally’ due to restricted gene exchange with other anatomical areas, producing distinctive sequences within each site [32]. It could also occur if HIV provirus-harboring cells undergo differential clonal expansion dynamics across sites, producing populations that differ from one another in terms of their relative frequencies of clonally-expanded sequences [33, 34]. Early studies uncovered differences in antiretroviral drug resistance profiles in lung and blood-derived sequences [35], and evidence of genetically compartmentalized HIV populations in blood and lung [36, 37], but these studies were not undertaken in context of suppressive ART.

A recent study of proviral diversity in tissues obtained at autopsy from ART-suppressed individuals, including three from whom lung tissue was sampled, revealed genetically diverse proviruses in lung [38], though no formal compartmentalization tests were applied. Another study of 18 individuals reported limited blood-lung compartmentalization, but 15 (83%) of the participants were viremic at the time of specimen collection, such that sampled proviruses would have largely represented actively-replicating, rather than archived viral strains [39]. A recent study of diverse tissues obtained at autopsy after long-term ART revealed clonally-expanded cell populations across multiple tissues with limited overall genetic compartmentalization, but lung-derived sequences were only available for two participants [40]. Another recent study analyzed *env* and *nef* sequences from autopsy tissues of five ART-suppressed individuals, three for whom lung-derived HIV sequences were obtained, but blood was not analyzed [41]. No studies to our knowledge have investigated blood/lung proviral diversity on long-term ART in context of HIV RNA populations replicating in plasma prior to ART initiation.

To extend our understanding of blood/lung proviral compartmentalization during long-term ART, we isolated >800 subgenomic proviral sequences by single-genome approaches from blood and bronchoalveolar lavage (BAL) from nine individuals. For two participants, we also interpret on-ART proviral diversity in context of HIV’s within-host evolutionary history, using HIV RNA sequences from plasma collected up to 18 years prior to proviral sampling.

## Results

### HIV sequence characterization in blood, lung, and plasma

At the time of blood and lung sampling, all nine participants had been virologically suppressed on ART for a median of 9 (IQR, 4-13) years (Table 1). In total, we isolated 1,025 subgenomic proviral sequences (*nef* region) by single-genome amplification from blood (N = 788) and lung (N = 237). After removal of 138 defective or hypermutated sequences, and five putative within-host recombinants, 882 intact *nef* proviral sequences remained: 699 (median 78; IQR, 58-90 per participant) from blood and 183 (median 14, IQR 6-37 per participant) from lung. For participants 4 and 6, we isolated an additional 91 and 211 HIV RNA *nef* sequences, respectively, by single-genome amplification from longitudinal pre-ART plasma samples collected up to 18 years prior to proviral sampling.

**Table 1.** Participant information and HIV sampling details.

| ID | Sex | Years of ART prior to proviral sampling | Intact proviral lung sequences<br>Total N (distinct n; %) | Intact proviral blood sequences<br>Total N (distinct n; %) | pre-ART plasma HIV RNA sequences<sup>a</sup><br>Total N (distinct n; %) |
| --- | --- | --- | --- | --- | --- |
| 1 | M | 8 | 29 (29; 100%) | 88 (64; 72%) | - |
| 2 | M | 15 | 31 (13; 42%) | 60 (50; 83%) | - |
| 3 | M | 0.6 | 6 (6; 100%) | 75 (60; 80%) | - |
| 4 | M | 18 | 42 (7; 17%) | 78 (53; 68%) | 91 (65; 71%) |
| 5 | M | 5 | 6 (6; 100%) | 91 (35; 38%) | - |
| 6 | M | 12 | 43 (18; 42%) | 126 (56; 44%) | 211 (99; 47%) |
| 7 | M | 10 | 4 (4; 100%) | 79 (57; 72%) | - |
| 8 | F | 9 | 14 (2; 14%) | 47 (44; 91%) | - |
| 9 | M | 3 | 8 (7; 88%) | 55 (40; 73%) | - |
| **Total** |  |  | **183 (92; 50%)** | **699 (459; 66%)** | **302 (164; 54%)** |

<sup>a</sup>Plasma HIV sequences were collected from three pre-ART timepoints from each of participants 4 and 6. Pre-ART plasma was not available for the other participants.

Participants differed markedly in terms of their overall number and anatomical distribution of distinct proviral *nef* sequences (Table 1 and Fig 1A). Participant 1 for example continued to frequently yield distinct sequences even after >100 proviruses had been collected, indicating that we did not capture the full extent of their diversity (Fig 1A). By contrast, more than 50% of participant 6’s sequences were duplicates. Overall, there was no consistent pattern in the frequency of distinct sequences recovered from blood versus lung (Wilcoxon matched pairs signed rank test p > 0.99; Fig 1B). This suggests that one compartment is not inherently more likely to harbor clonally-expanded populations than the other, though it is interesting that the lungs of two participants (4 and 8) were nearly entirely comprised of identical sequences, while no such phenomenon was observed in the blood of any participant. A phylogenetic tree inferred from all intact HIV sequences confirmed that each participant formed a monophyletic clade with branch support values of 100%, and all participants harbored HIV subtype B (Fig 1C). Overall, participant 3 exhibited the highest within-host diversity (calculated as the mean patristic or “tip-to-tip” distance between all pairs of distinct sequences) whereas participant 9 exhibited the least diversity (Fig 1C, inset). Although clinical histories are not available for all participants, these observations are consistent with participant 3 having had HIV for at least 15 years prior to suppressing viremia on ART, whereas participant 9 initiated ART shortly following diagnosis (early ART limits HIV evolution and diversity [42, 43]).

**Fig 1.**
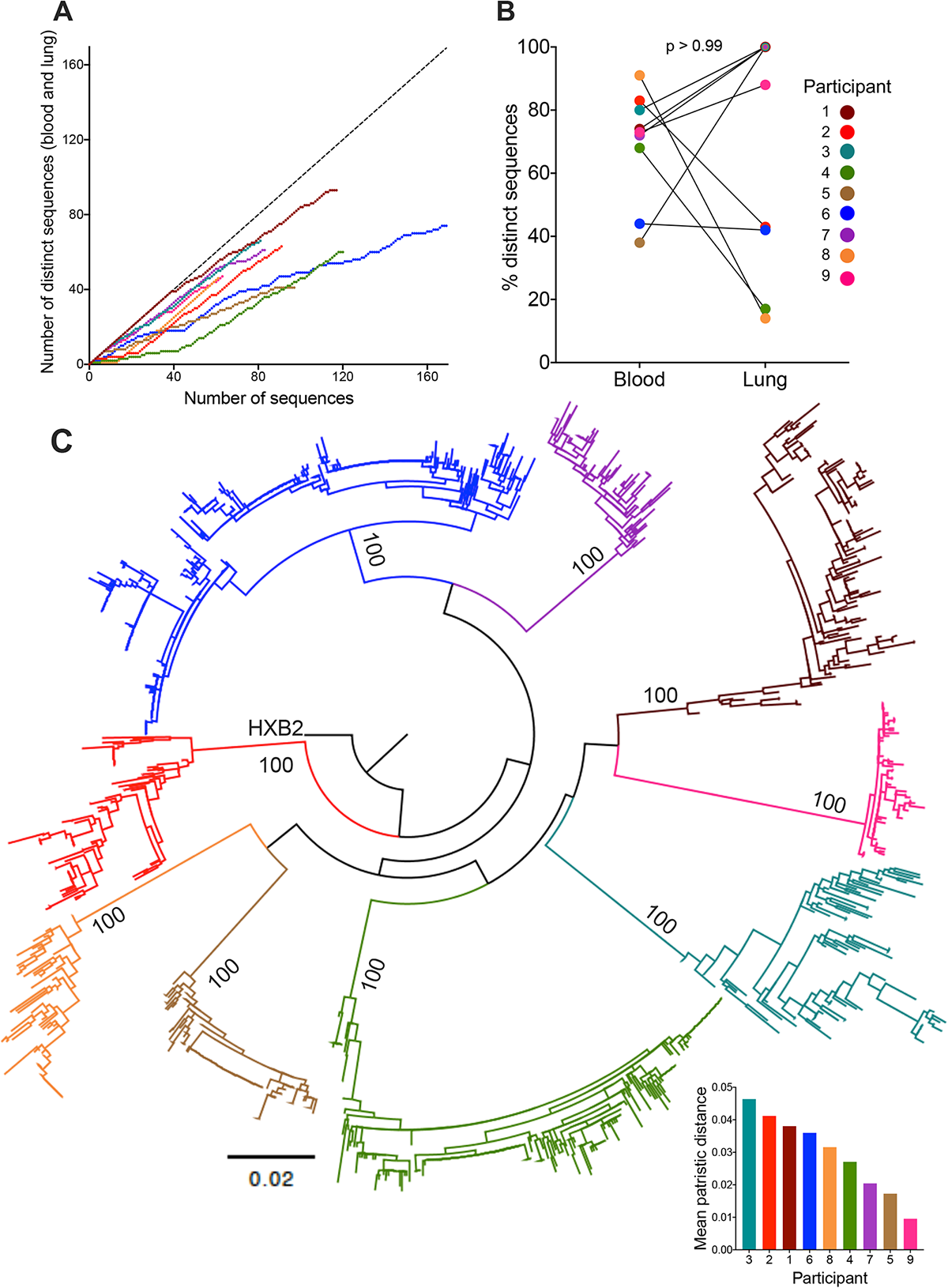
Within-host HIV sequence characterization. (A) Total number of distinct HIV sequences collected from blood and lung per participant, as a function of their overall number of sequences collected. This helps estimate the extent to which sampled sequences captured within-host diversity. (B) Proportion of distinct proviral sequences isolated from blood and lung of each participant; p-value calculated by Wilcoxon matched pairs signed rank test. (C) Maximum-likelihood phylogeny inferred from 882 proviral and 302 plasma HIV RNA sequences isolated from participants, rooted on the HIV subtype B reference strain HXB2. Numbers on internal branches indicate bootstrap values. Scale in estimated substitutions per nucleotide site. Inset: participants ranked from highest to lowest within-host genetic diversity (calculated as mean patristic distances of distinct sequences).

### Proviral diversity and compartmentalization in blood and lung

We next investigated proviral diversity and compartmentalization in blood and lung, beginning with participant 3 (whose proviruses were the most diverse) and ending with participant 9 (whose proviruses were the least diverse) (Figs 2 to 6). For each participant, the highest likelihood phylogeny from among 7,500 within-host reconstructions is shown. As compartmentalization tests assess different genetic features of the populations in question, it is standard to apply more than one type of test for reliable compartmentalization inference [44]. We applied four tests: two that were genetic distance-based and two that were tree-based and defined a *priori* that a dataset would be declared compartmentalized if at least two tests returned statistically significant results (i.e., p < 0.05).

**Fig 2.**
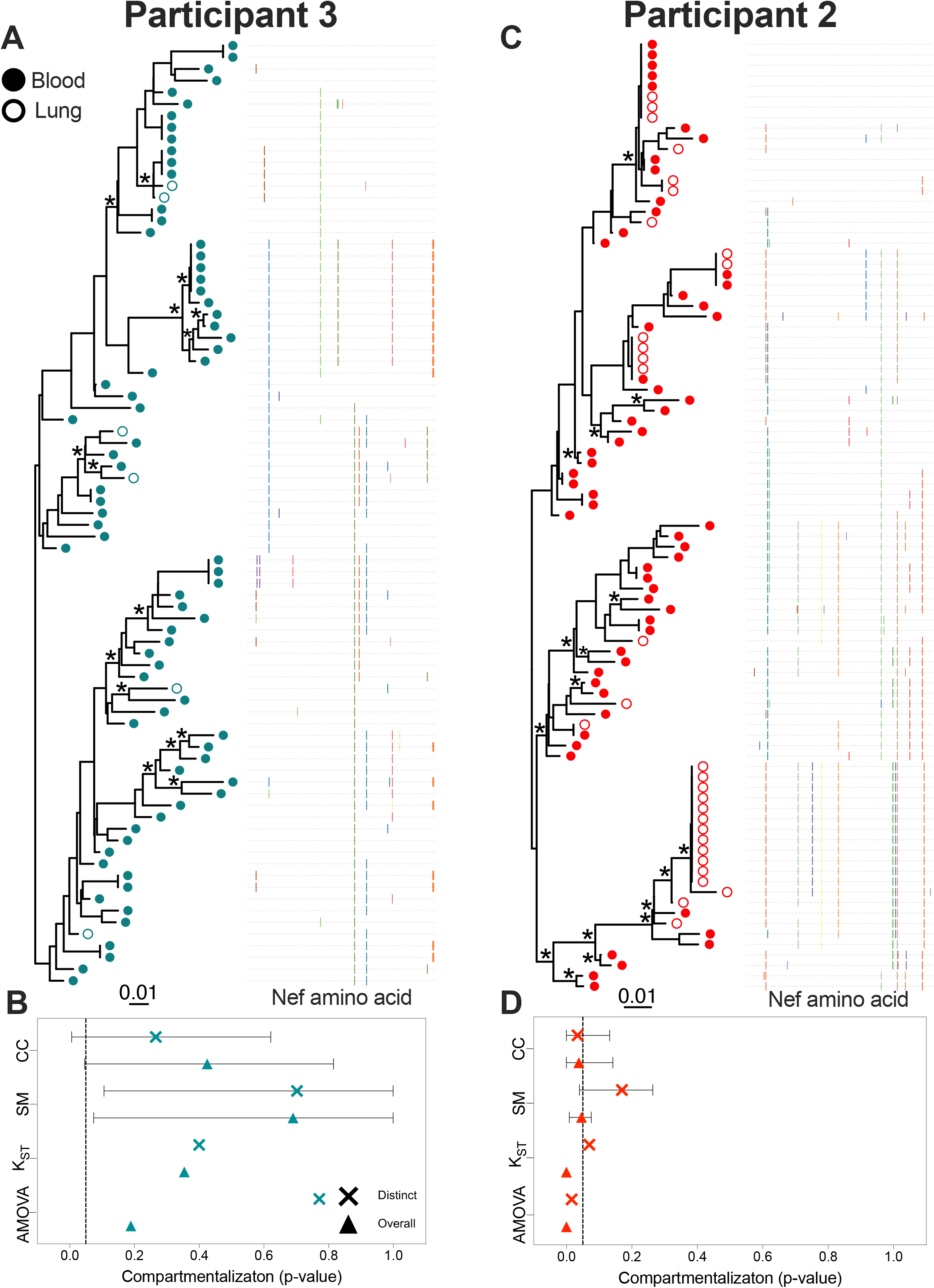
Within-host phylogenies and compartmentalization results for participants 3 and 2. Each participant’s top panel shows the highest likelihood phylogeny derived from Bayesian inference, rooted at the midpoint. Filled circles denote blood sequences; open circles denote lung sequences. Asterisks identify nodes supported by posterior probabilities ≥70%. The adjacent plot shows amino acid diversity of each sequence, with coloured ticks denoting non-synonymous substitutions with respect to the sequence at the top of the phylogeny. The bottom panel shows p-values of the four tests of genetic compartmentalization between blood and lung (AMOVA = Analysis of Molecular Variance; KST = Hudson, Boos and Kaplan’s nonparametric test for population structure; SM = Slatkin-Maddison test; CC = Correlation Coefficients test). The “×” symbol shows the p-value based on distinct sequences per compartment, and the triangle shows the p-value based on overall sequences. For SM and CC tests, the bars represent the 95% HPD interval of the p-value derived from all 7,500 trees.

The genetic distance-based tests were Analysis of Molecular Variance (AMOVA) [45], and Hudson, Boos and Kaplan’s nonparametric test for population structure (KST) [46]. AMOVA calculates an association based on the genetic diversity of the sequences between and within compartments, where variability is calculated from the sum of the squared genetic distances between sequences. KST compares the mean pairwise distances between sequences from different versus the same compartment. Compartmentalization is supported if the latter are smaller than the former, with statistical significance derived via a population-structure randomization test. Both test statistics (phi [ϕ] and KST, respectively) range from 0 (no compartmentalization) to 1 (complete compartmentalization), where p-values are derived from 1000 permutations (Table 2). The two tree-based methods were the Slatkin-Maddison (SM) test [47] and the Correlation Coefficients (CC) test [48]. SM determines the minimum number of migrations between compartments to explain the distribution of compartments on the tree tips: the smaller the number of migrations, the stronger the compartmentalization support. Statistical support is based on the number of migrations that would be expected in a randomly-structured population, simulated by permuting compartment labels between sequences. The CC test correlates the phylogenetic distances between two sequences (defined here as the number of internal branches separating them in the tree) with information on their compartment of isolation. Coefficients can range from -1 to +1, where larger positive values denote stronger support for compartmentalization, and values closer to zero (and negative values) denote no compartmentalization. Statistical support is derived from estimating the distribution of these coefficients by permuting compartment labels between sequences. The tree-based tests were applied to all 7,500 within-host phylogenies per participant, and p-values averaged. Each dataset was analyzed two ways: first, restricting to only distinct sequences *per compartment* (“distinct” analysis; where any sequence present in both lung and blood was represented once per compartment), and second, including all sequences (“overall” analysis). This is because identical HIV sequences, especially when differentially abundant across compartments, increase the likelihood of obtaining a significant compartmentalization result, but such distributions are likely the result of differential clonal expansion rather than divergent HIV evolution between compartments [34, 39].

**Table 2.**
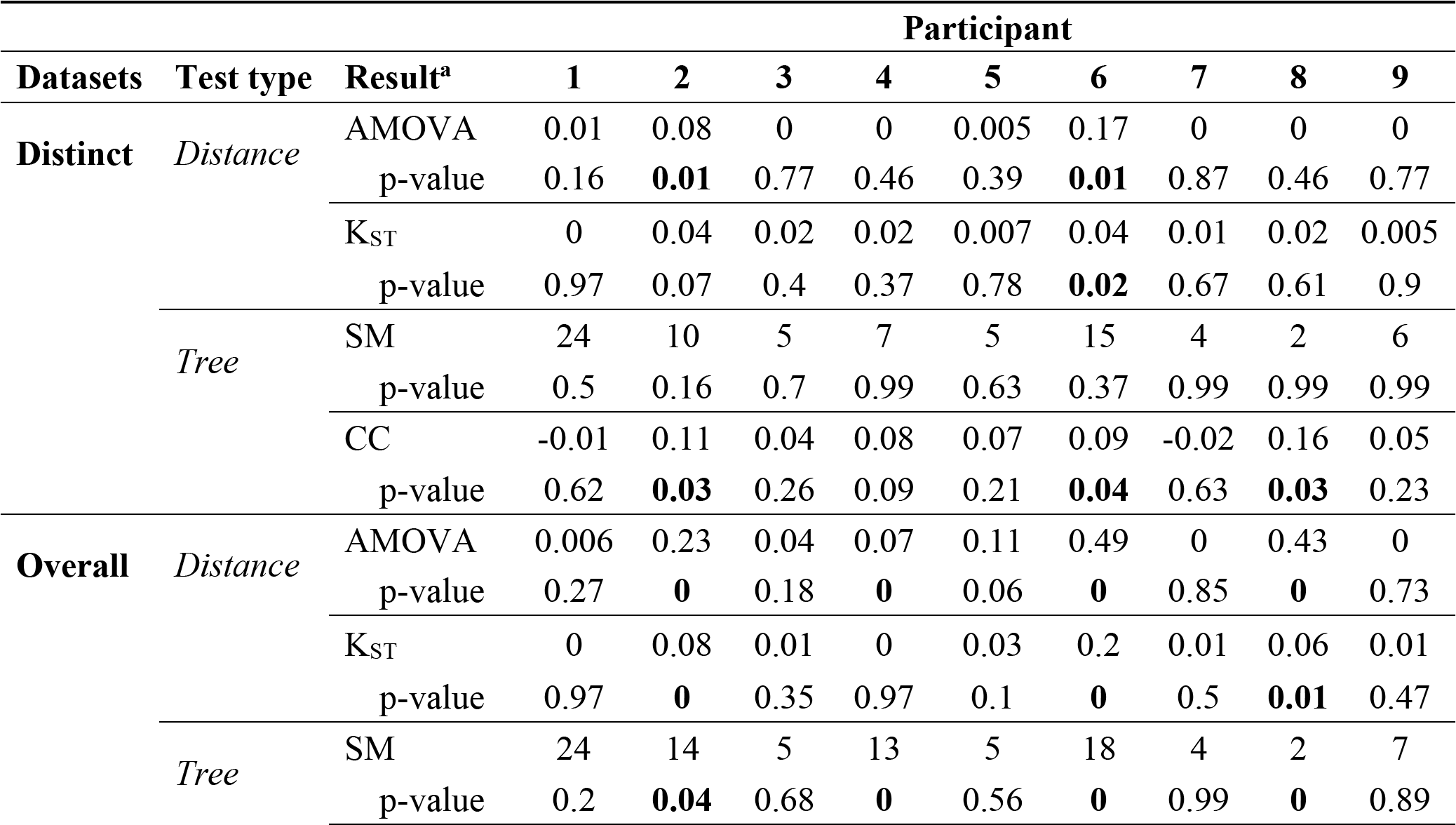
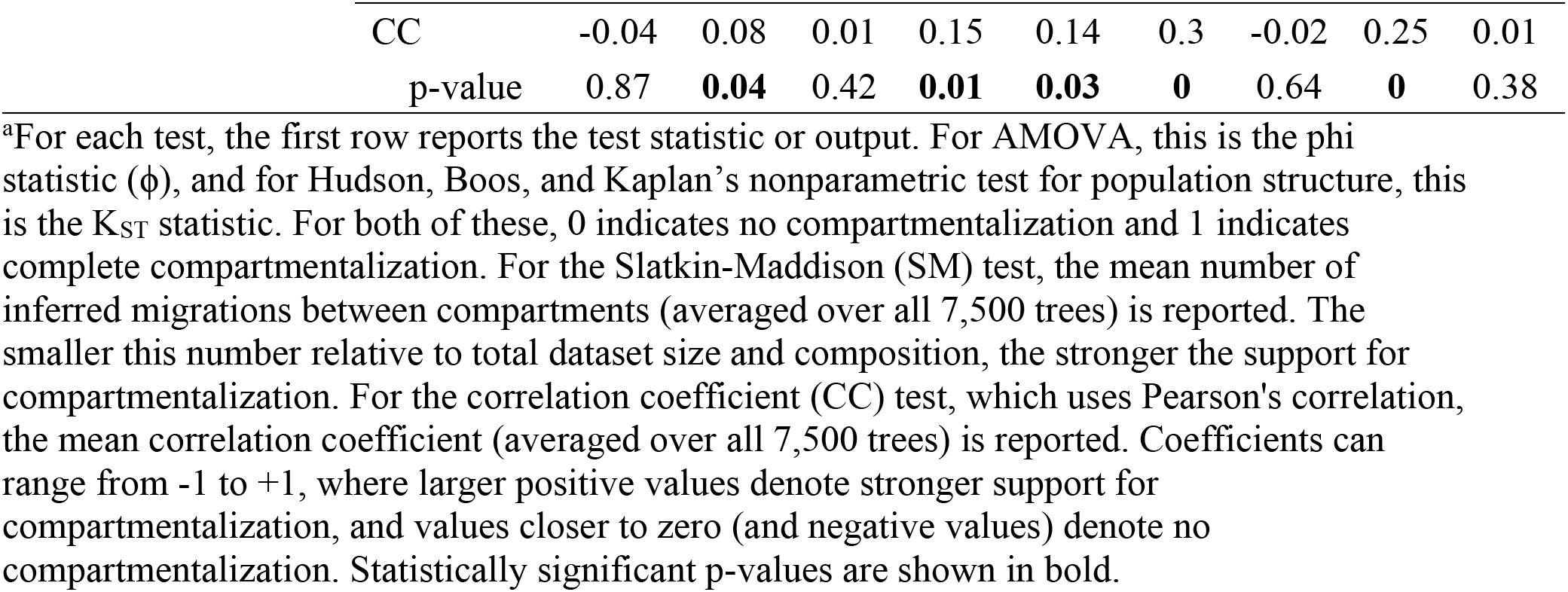
Blood-lung genetic compartmentalization results.

From participant 3, we recovered 75 *nef* sequences from blood (80% distinct), the most abundant of which was recovered five times (Fig 2A). Although we recovered only six sequences from lung, all were unique, quite divergent from one another (see how they intersperse throughout the within-host phylogeny) and did not match any sequence in blood. The highlighter plot adjacent to the phylogeny illustrates within-host amino acid diversity. For participant 3, none of the four tests supported compartmentalization between blood and lung regardless of whether identical sequences were included in the analysis or not (Fig 2B).

From participant 2 we recovered 60 sequences from blood (83% distinct in this compartment) and 31 from lung (42% distinct in this compartment) (Fig 2C). Four sequences were found in both blood and lung; in three of these cases there were two or more copies of this sequence in one or both compartments. The most abundant sequence was observed 12 times (all in lung). Our *a priori* definition for compartmentalization required statistically significant evidence in at least two of the four tests. By this definition, participant 2 displayed modest yet statistically significant evidence for compartmentalization when considering only distinct sequences per compartment (AMOVA ϕ = 0.08, p = 0.01; CC coefficient = 0.11, p = 0.03), though in both cases the test statistic value was low (Fig 2D and Table 2). When all sequences were considered, all four tests returned significant results (with AMOVA and KST yielding p = 0, and SM and CC yielding p = 0.04), though the overall magnitude of compartmentalization remained modest (e.g. AMOVA ϕ increased from 0.08 to 0.23, while the CC value decreased from 0.11 to 0.08).

From participant 1 we recovered 88 sequences from blood (72% distinct in this compartment) and 29 from lung (100% distinct in this compartment), though seven sequences were found in both compartments. The most abundant sequence was observed six times (five in blood, one in lung) (Fig 3A). No evidence of compartmentalization was detected by any test regardless of the inclusion of identical sequences (Fig 3B).

**Fig 3.**
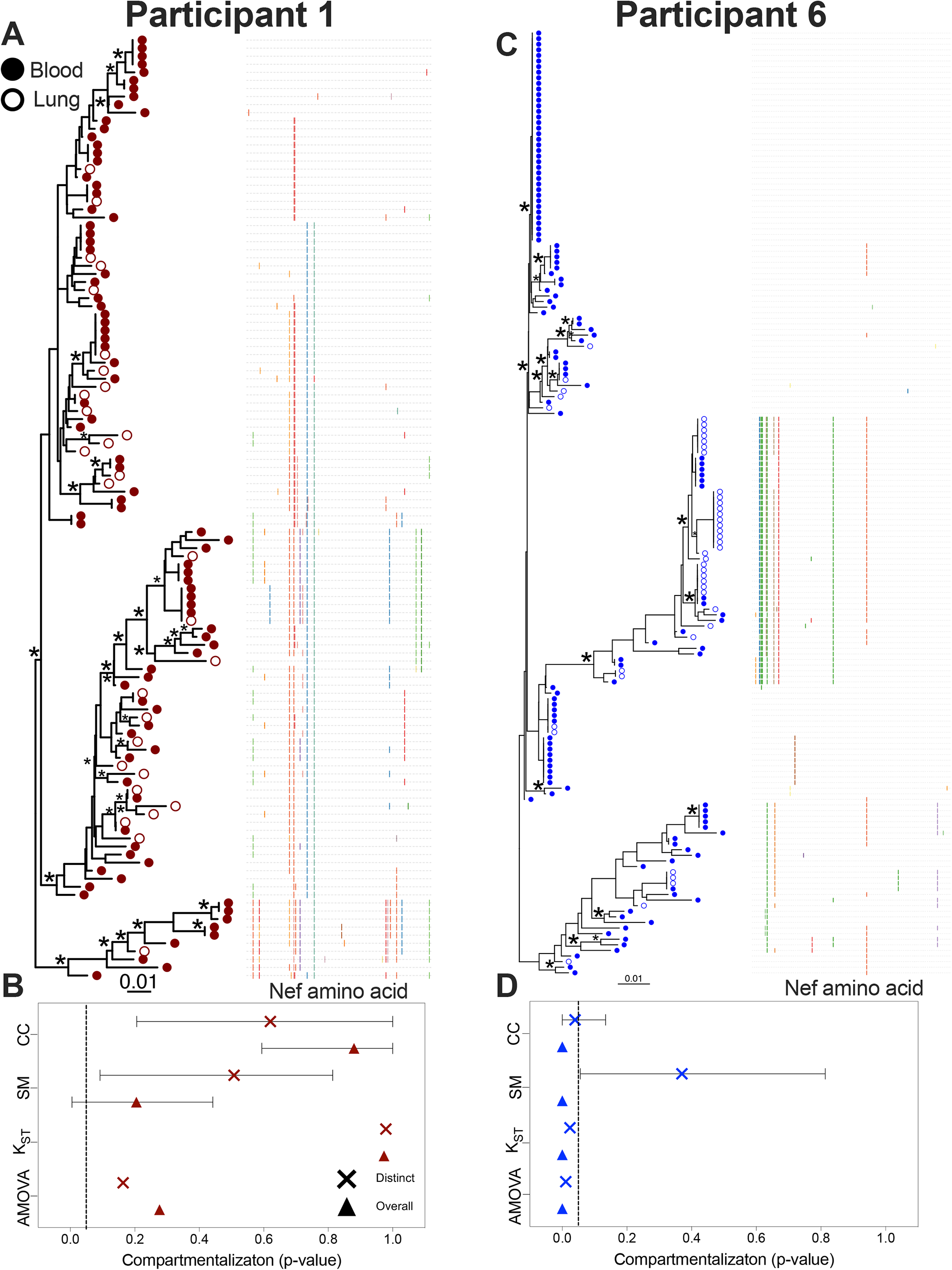
Within-host phylogenies and compartmentalization results for participants 1 and 6. Each participant’s top panel shows the highest likelihood phylogeny derived from Bayesian inference, rooted at the midpoint. Filled circles denote blood sequences; open circles denote lung sequences. Asterisks identify nodes supported by posterior probabilities ≥70%. The adjacent plot shows amino acid diversity of each sequence, with coloured ticks denoting non-synonymous substitutions with respect to the sequence at the top of the phylogeny. The bottom panel shows p-values of the four tests of genetic compartmentalization between blood and lung (AMOVA = Analysis of Molecular Variance; KST = Hudson, Boos and Kaplan’s nonparametric test for population structure; SM = Slatkin-Maddison test; CC = Correlation Coefficients test). The “×” symbol shows the p-value based on distinct sequences per compartment, and the triangle shows the p-value based on overall sequences. For SM and CC tests, the bars represent the 95% HPD interval of the p-value derived from all 7,500 trees.

Participant 6 was distinctive in that they harbored the highest overall frequency of identical sequences (only 44% of the 126 sequences recovered from blood and 42% of those from lung were distinct in those compartments) and that these identical sequences tended to be present in a single compartment only (Fig 3C). The most abundant sequence was observed 38 times (all in blood). An additional eight sequences were observed five or more times, most often in a single compartment, though there were four instances where a sequence was observed in both blood and lung. Like participant 2, participant 6’s dataset exhibited modest yet statistically significant evidence of compartmentalization when considering only distinct sequences per compartment (AMOVA ϕ = 0.17, p = 0.01; KST = 0.04, p = 0.02; CC coefficient = -0.02, p = 0.04). Support for compartmentalization became more pronounced when all sequences were considered (with AMOVA ϕ for example reaching 0.49, and all tests returning p = 0) (Fig 3D and Table 2).

Participant 8 was remarkable in that 13 out of 14 sequences isolated from lung were identical and distinct to this compartment, whereas blood sequences were relatively diverse with few duplicates (Fig 4A). All four tests returned significant evidence of compartmentalization when considering all sequences (with AMOVA, SM and CC returning p = 0, and KST returning p = 0.01), but not when considering only distinct sequences (Fig 4B).

**Fig 4.**
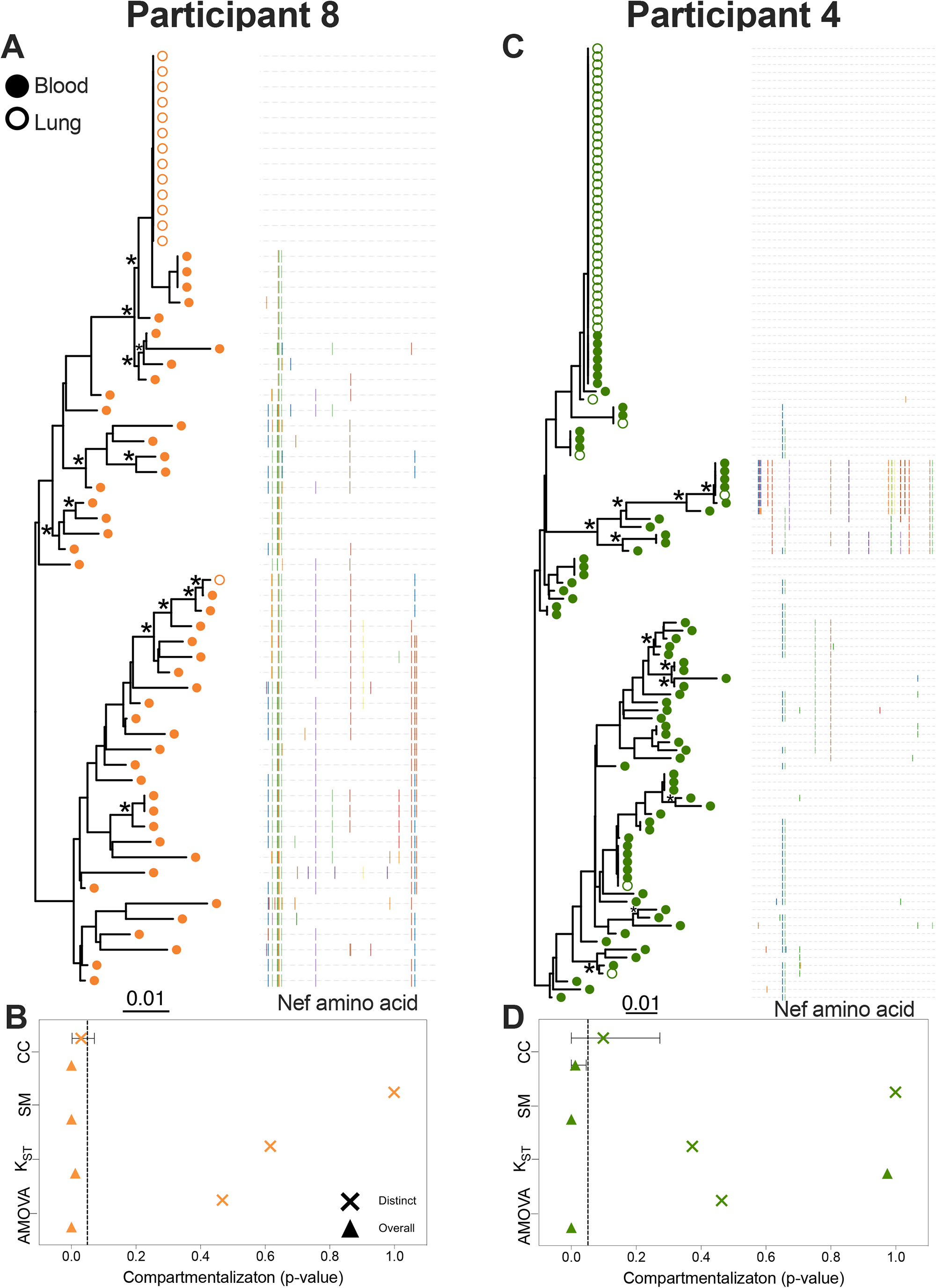
Within-host phylogenies and compartmentalization results for participants 8 and 4. Each participant’s top panel shows the highest likelihood phylogeny derived from Bayesian inference, rooted at the midpoint. Filled circles denote blood sequences; open circles denote lung sequences. Asterisks identify nodes supported by posterior probabilities ≥70%. The adjacent plot shows amino acid diversity of each sequence, with coloured ticks denoting non-synonymous substitutions with respect to the sequence at the top of the phylogeny. The bottom panel shows p-values of the four tests of genetic compartmentalization between blood and lung (AMOVA = Analysis of Molecular Variance; KST = Hudson, Boos and Kaplan’s nonparametric test for population structure; SM = Slatkin-Maddison test; CC = Correlation Coefficients test). The “×” symbol shows the p-value based on distinct sequences per compartment, and the triangle shows the p-value based on overall sequences. For SM and CC tests, the bars represent the 95% HPD interval of the p-value derived from all 7,500 trees.

Participant 4 was remarkable in that they harbored a single sequence that was recovered 42 times (36 times in lung, 7 in blood) (Fig 4C). An additional four sequences were shared across compartments, but otherwise the remaining blood and lung sequences were relatively diverse. There was no evidence of compartmentalization when considering distinct sequences only, but three of the four tests returned significant results when considering all sequences (AMOVA and SM returned p = 0; CC returned p = 0.01) (Fig 4D).

For participants 7, 5 and 9, we recovered only 4, 6 and 8, lung sequences, respectively (Figs 5A and C, 6A). Despite this limited sampling, lung sequences were relatively divergent from one another, and most were distinct. In all three participants however, at least one lung sequence was also found in blood. Participant 5 was also remarkable for the recovery of 41 identical sequences, all in blood. For these three participants, no test identified significant compartmentalization regardless of the inclusion of identical sequences (Figs 5B and D, 6B).

**Fig 5.**
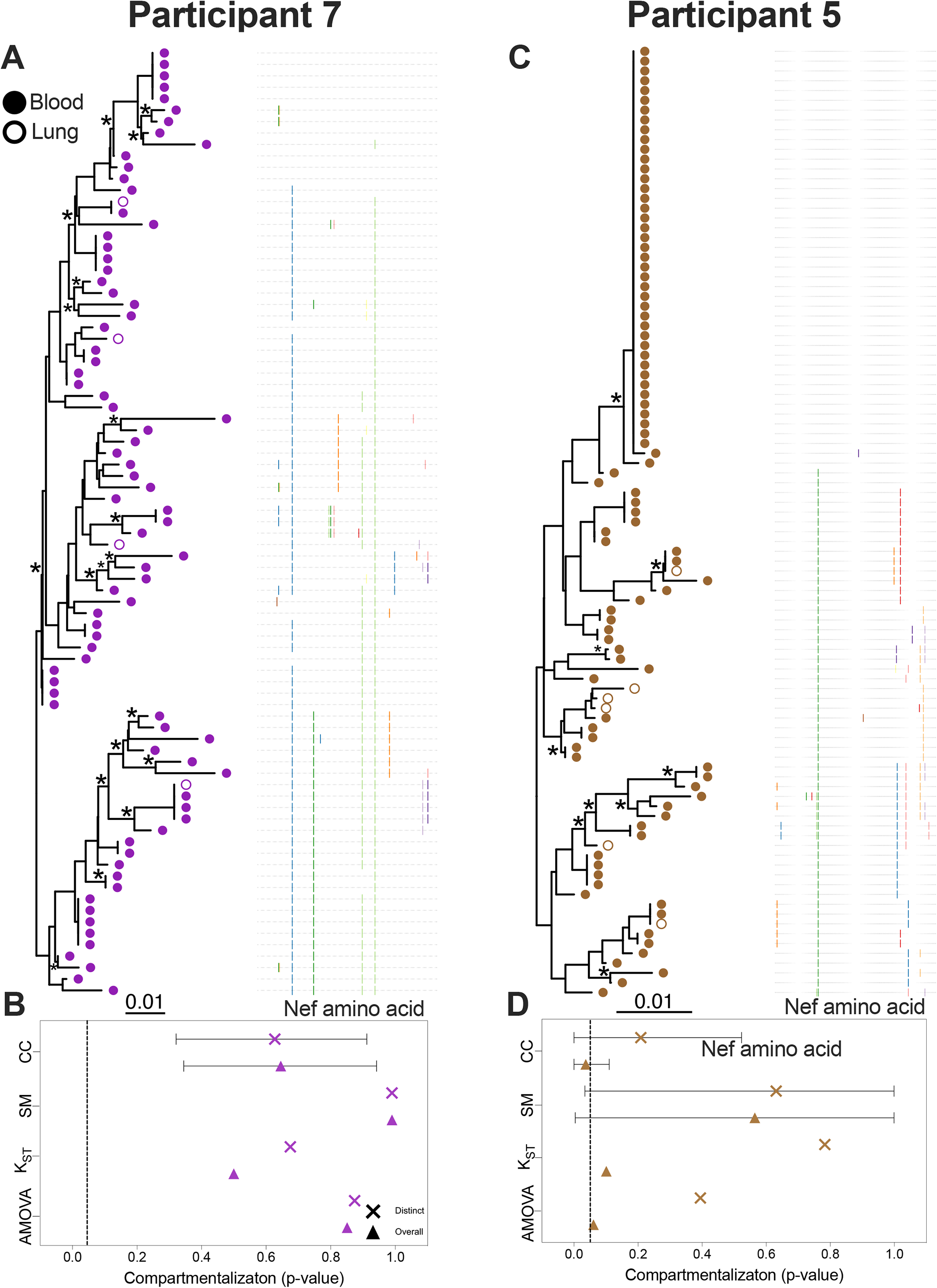
Within-host phylogenies and compartmentalization results for participants 7 and 5. Each participant’s top panel shows the highest likelihood phylogeny derived from Bayesian inference, rooted at the midpoint. Filled circles denote blood sequences; open circles denote lung sequences. Asterisks identify nodes supported by posterior probabilities ≥70%. The adjacent plot shows amino acid diversity of each sequence, with coloured ticks denoting non-synonymous substitutions with respect to the sequence at the top of the phylogeny. The bottom panel shows p-values of the four tests of genetic compartmentalization between blood and lung (AMOVA = Analysis of Molecular Variance; KST = Hudson, Boos and Kaplan’s nonparametric test for population structure; SM = Slatkin-Maddison test; CC = Correlation Coefficients test). The “×” symbol shows the p-value based on distinct sequences per compartment, and the triangle shows the p-value based on overall sequences. For SM and CC tests, the bars represent the 95% HPD interval of the p-value derived from all 7,500 trees.

**Fig 6.**
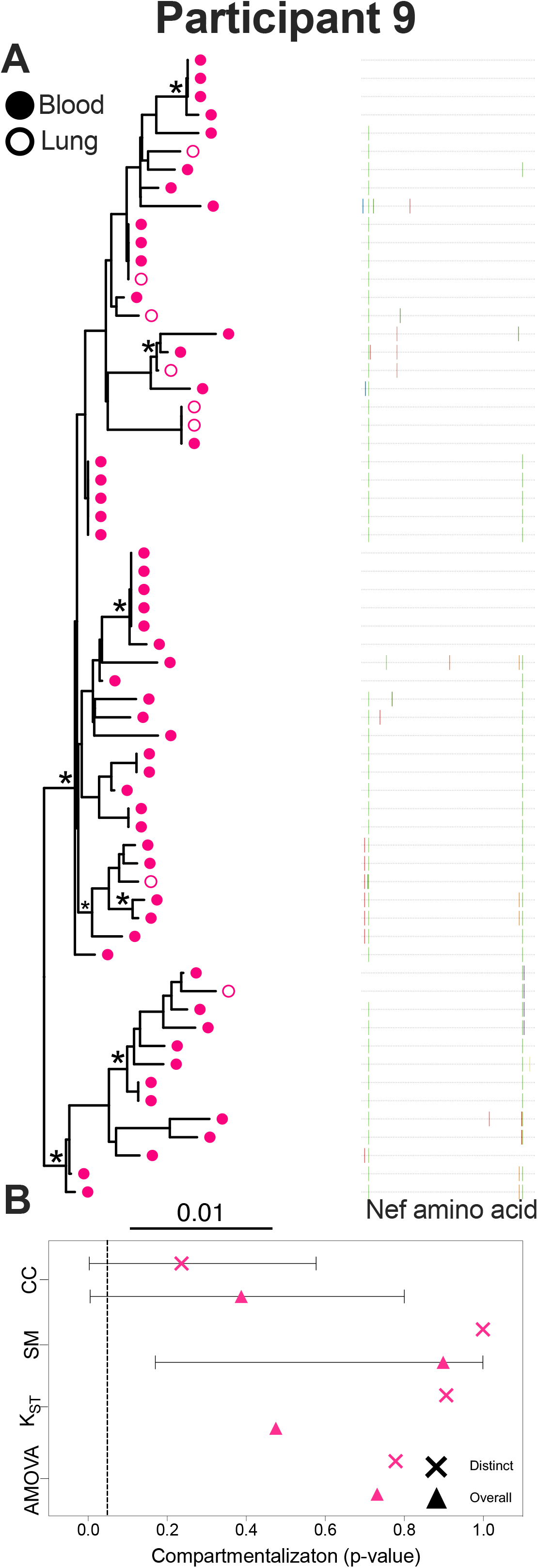
Within-host phylogeny and compartmentalization results for participant 9. Each participant’s top panel shows the highest likelihood phylogeny derived from Bayesian inference, rooted at the midpoint. Filled circles denote blood sequences; open circles denote lung sequences. Asterisks identify nodes supported by posterior probabilities ≥70%. The adjacent plot shows amino acid diversity of each sequence, with coloured ticks denoting non-synonymous substitutions with respect to the sequence at the top of the phylogeny. The bottom panel shows p-values of the four tests of genetic compartmentalization between blood and lung (AMOVA = Analysis of Molecular Variance; KST = Hudson, Boos and Kaplan’s nonparametric test for population structure; SM = Slatkin-Maddison test; CC = Correlation Coefficients test). The “×” symbol shows the p-value based on distinct sequences per compartment, and the triangle shows the p-value based on overall sequences. For SM and CC tests, the bars represent the 95% HPD interval of the p-value derived from all 7,500 trees.

To summarize, when considering only distinct sequences per compartment, we detected statistically significant support for blood-lung genetic compartmentalization in only two of nine individuals studied (participants 2 and 6). When considering all within-host sequences, this number increased to four (participants 2, 4, 6 and 8). By contrast, five of the nine participants (1, 3, 5, 7 and 9) exhibited no significant compartmentalization by any analysis (Fig 7A).

**Fig 7.**
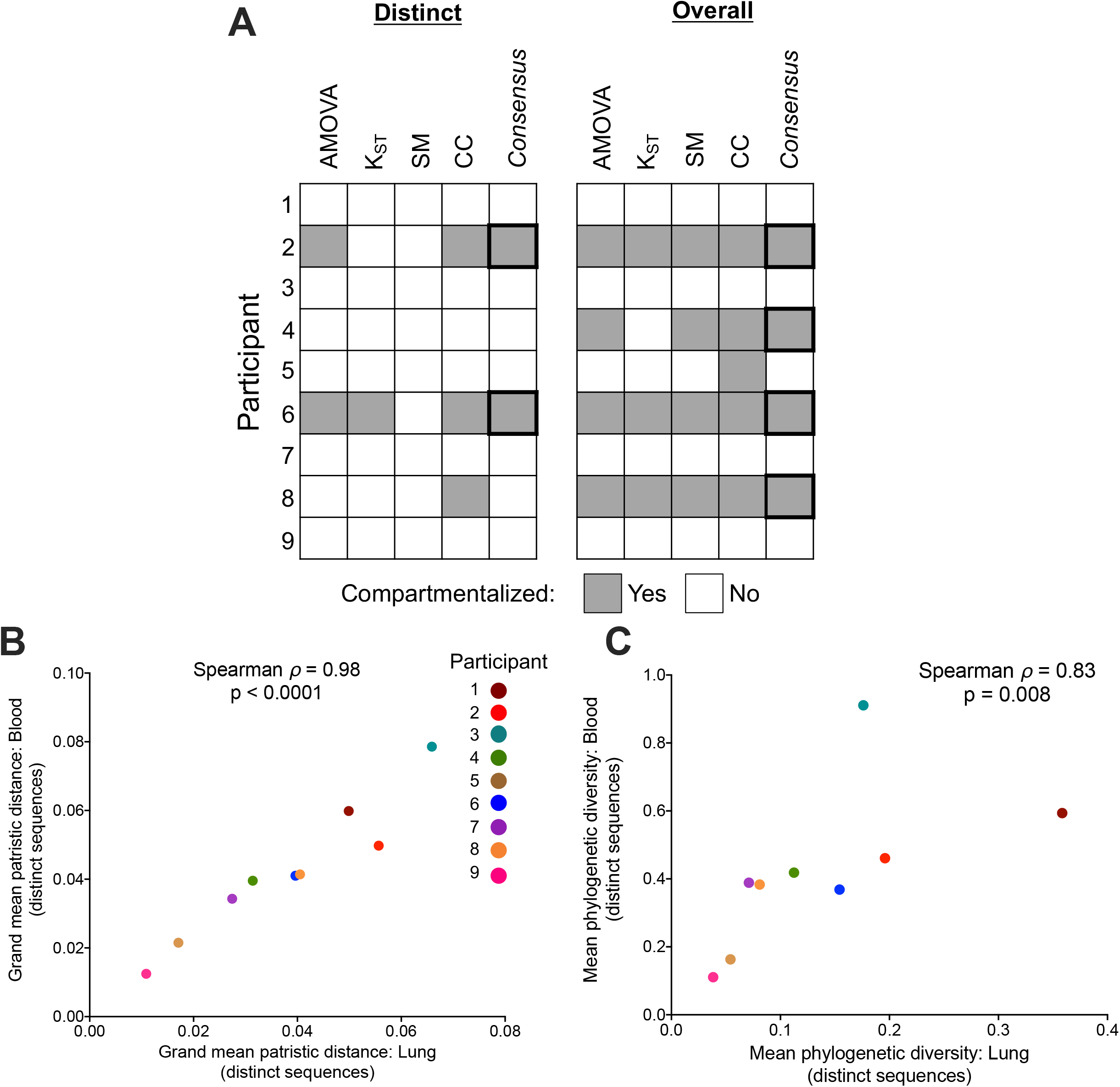
Summary of genetic compartmentalization results and proviral diversity. (A) Summary of genetic compartmentalization results for the four tests used, along with the “consensus” result, where a dataset is declared compartmentalized if at least two tests gave statistically significant results. Grey squares denote compartmentalization; white squares denote no compartmentalization. The “Distinct” panel summarizes compartmentalization results when limiting analysis to distinct sequences per compartment; the “Overall” panel summarizes results when all sequences are included. (B) Spearman’s correlation between blood and lung proviral diversity, where values represent the grand mean of within-host patristic distances separating all pairs of distinct sequences per compartment, averaged over all 7,500 trees per participant. (C) Spearman’s correlation between blood and lung proviral diversity, where values represent mean phylogenetic diversity of distinct sequences per compartment, computed from all 7,500 trees per participant.

Despite major inter-individual heterogeneity in proviral diversity and composition, one notable observation nevertheless emerged: within-host diversity of distinct proviral sequences recovered from blood correlated significantly with the diversity of distinct sequences recovered from lung (Figs 7B and C). Here, we calculated within-host diversity two ways: first by computing the mean patristic (tip-to-tip) distances separating blood and lung sequences from one another in all 7,500 phylogenetic reconstructions per participant and then averaging these values across all trees, and second by computing the overall mean phylogenetic diversity per compartment (as the sum of edge lengths of distinct sequences per compartment) averaged over all trees. The first measure of diversity yielded a Spearman ρ = 0.98, p < 0.0001 between blood and lung (Fig 7B) while the second yielded a Spearman ρ = 0.83, p = 0.008 (Fig 7C).

### Exploring proviral diversity in context of pre-ART viral populations

For two participants, 4 and 6, the availability of pre-ART plasma samples collected up to 18 years prior to proviral sampling allowed us to contextualize the diversity of proviruses persisting on ART with that of HIV populations that previously replicated in plasma, revealing insights into within-host proviral dynamics and longevity.

Participant 6 was diagnosed with HIV in June 1982, and first initiated ART in 1997 (Fig 8A). Treatment was interrupted from February to April 2002, and again from February 2006 to August 2009, during which time pVL increased to ≥4 log10. There was another brief viremic episode in 2012 (average 215 RNA copies/mL) after which suppression was re-established. A pre-ART plasma sample from October 1996, along with samples from 2006 and 2009 during the second treatment interruption, were available for study. We isolated 211 intact plasma HIV RNA sequences (68 from 1996; 86 from 2006; 57 from 2009), and inferred 7,500 phylogenies from these along with the 169 proviral sequences isolated from blood and lung during ART.

**Fig 8.**
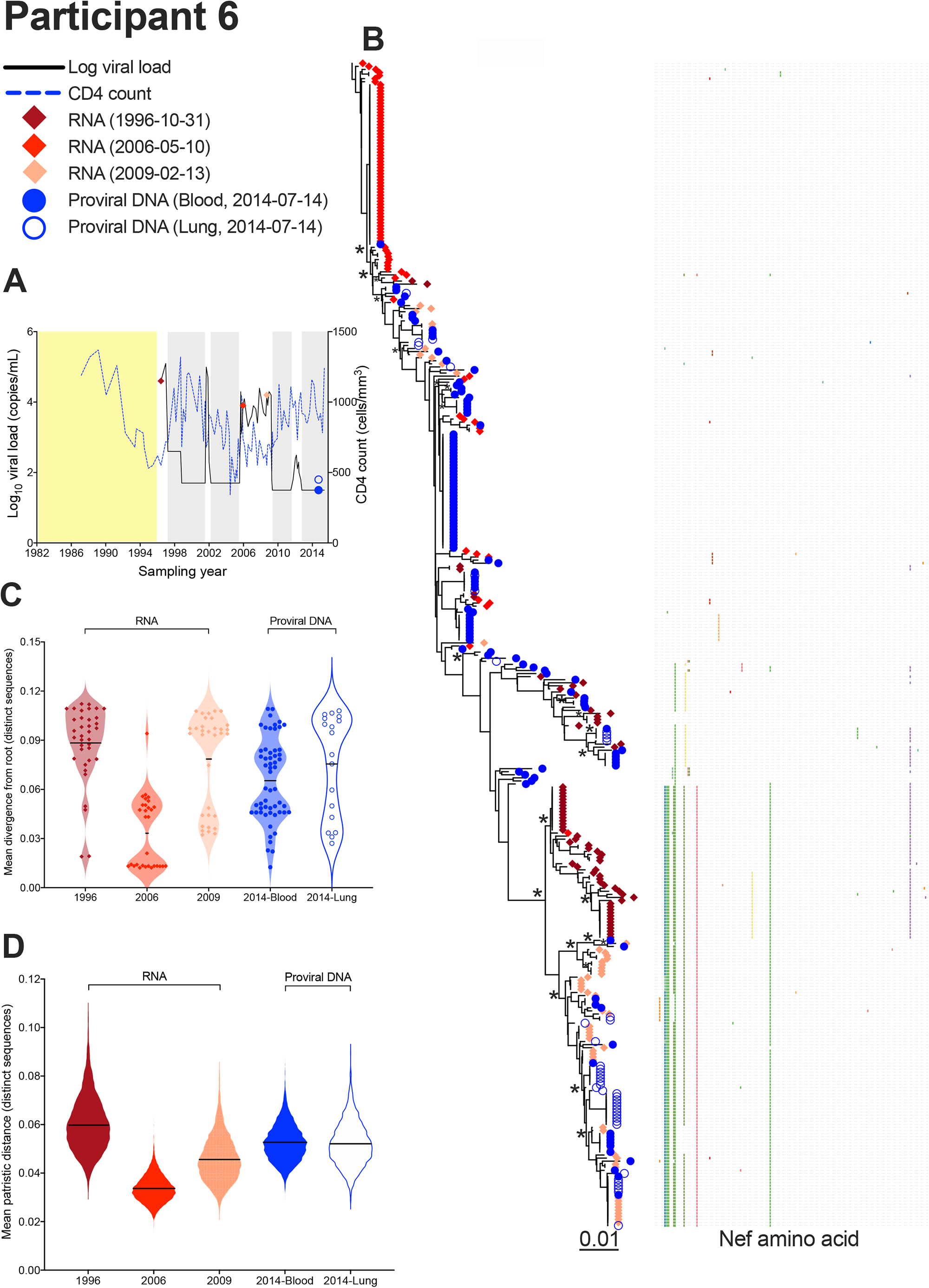
Interpreting proviral diversity in context of prior within-host HIV evolution: participant 6. (A) Plasma viral load (solid black line), CD4^+^ T-cell count (dashed blue line) and samples analyzed (colored diamonds and circles). Yellow shading denotes period following infection where clinical information is incomplete. Grey shading denotes periods of suppressive ART as per the HIV RNA pVL assay used at the time (until 1998, values <400 copies/mL indicated undetectable viremia; later assays used a <50 copies/mL cutoff) [81]. (B) Highest likelihood phylogeny derived from Bayesian inference, rooted as described in the text. Scale in estimated substitutions per nucleotide site. The adjacent plot shows amino acid diversity, with coloured ticks denoting non-synonymous substitutions with respect to the sequence at the top of the tree. Asterisks identify nodes supported by posterior probabilities ≥70%. (C) Mean divergence from the root of each distinct sequence, averaged over all 7,493 trees, stratified by sampling date and sample type. The black line represents the grand mean. (D) Within-host HIV sequence diversity, calculated as the mean patristic distance between all distinct sequences collected at that time point for that sample, where the values for each of the 7,493 trees are represented as a violin plot. The black line represents the grand mean.

Interpreting on-ART proviral diversity in context of HIV’s within-host evolutionary history ideally requires rooting the tree at the most recent common ancestor (MRCA) of the collected sequences. If the tree contains sequences from historic evolving within-host viral populations, this can be done by placing the root at the location that maximizes the correlation between the root-to-tip distances and the sampling dates of those evolving sequences [7, 49, 50]. This strategy works because evolving within-host HIV populations exhibit increasing divergence from the root over time. Rooting can be performed with as few as two time points, usually longitudinal pre-ART plasma samples, provided that sufficient evolution has occurred between them [7]. Longitudinal pre-ART plasma samples were not available for this participant, but uncontrolled HIV replication did occur between 2006 and 2009, where the 2006 sequences would have represented HIV variants newly emerging from the reservoir to repopulate plasma. We therefore rooted each of the participant’s 7,500 phylogenies at the location that maximized the correlation between root-to-tip distances and sampling time of plasma sequences collected in 2006 and 2009. The mean Pearson’s correlation coefficient supporting these root placements was 0.63 (95% HPD 0.56 to 0.70), where 7,493/7,500 (99.9%) trees returned p-values that reached our predefined cutoff for this analysis of p ≤ 0.001. A representative tree (the one with the highest likelihood) is shown in Fig 8B.

Assuming that the root represents the MRCA of the sequences collected, we can make some inferences about proviral dynamics. First, the plasma HIV RNA sequences that were circulating in 1996, just prior to ART initiation, ranged widely in terms of divergence from the MRCA (grand mean of 0.088 substitutions per nucleotide site; Fig 8C) and were also quite diverse (mean 0.059 substitutions per nucleotide site; Fig 8D). This is not unexpected given that these sequences represented the end products of nearly 15 years of uncontrolled within-host HIV evolution. The sequences emerging in plasma in 2006 by contrast were less divergent from the root than the 1996 ones (both the grand mean divergence from the root and diversity were a mean 0.033 substitutions per nucleotide site). This suggests that the replication-competent HIV variants that emerged from the reservoir to re-populate plasma in 2006 were from a relatively restricted subset that included variants that had circulated somewhat prior to 1996 (these are the sequences closest to the root, which have shorter root-to-tip distances than any 1996 sequence). These variants evolved over the next three years, as shown by the increased root-to-tip divergence and diversity (grand mean 0.078 and 0.045, respectively) of the 2009 plasma sequences compared to the 2006 ones. Interestingly, blood and lung proviral sequences sampled five years later during suppressive ART were most similar to the 2009 plasma HIV RNA population in terms of these metrics, with many proviral sequences clustering with 2009 plasma sequences (though clustering with 2006 and 1996 plasma sequences was also observed to some extent). Moreover, the average root-to-tip divergence and diversity of on-ART proviral sequences was lower than those of 1996 plasma sequences just prior to ART initiation. Overall, this suggests that participant 6’s proviral pool was substantially re-seeded during the 2006-2009 treatment interruption. The data also suggest that, while the sequences that originally rebounded in plasma in 2006 were ancestral to those that circulated at ART initiation, these were not *too* much more ancestral (as the root is not too much farther back than the least divergent pre-ART plasma sequence sampled in 1996). This relatively shallow root location, combined with the lack of proviral sequences close to this root, suggests that this individuals’ reservoir, when sampled in 2014, contained few or no sequences dating back to early infection. Of note, the 1996 plasma sequences featured a distinct subclade that was quite divergent from the root that contained only a single sequence from one other timepoint, a 2006 plasma sequence. This suggests that representatives of this clade, which were quite abundant in plasma just prior to ART initiation, did not persist in abundance in the reservoir.

Participant 4 was diagnosed with HIV in January 1987 but initiated ART only in February 2000 (Fig 9A). Viremia was generally suppressed thereafter except for brief episodes in 2004 and 2005. We isolated 36, 33 and 22 intact plasma HIV RNA sequences, respectively, from plasma sampled in January 2000 (pre-ART) and the 2004 and 2005 viremic episodes. As substantial HIV evolution could not have occurred between any of these time points, we rooted this participant’s 7,500 phylogenies using the HIV subtype B reference sequence HXB2 as an outgroup (representative tree in Fig 9B). Here, a very different picture emerges compared to participant 6. Notably, the proviruses sampled from participant 4 after almost 18 years on ART interspersed throughout the whole phylogeny and represented all sequences near the root, which was far deeper in the tree than any pre-ART HIV RNA sequence (Fig 9B). Indeed, on-ART proviruses ranged widely in terms of their root-to-tip distances but were nevertheless on average closer to the root than any plasma HIV populations sampled previously (e.g. lung mean proviral divergence was 0.049 substitutions per nucleotide site compared to 0.080 among sequences circulating in plasma just prior to ART; Fig 9C). Proviruses persisting in blood during ART were also on average more diverse (mean 0.051 substitutions per nucleotide site) than any plasma HIV populations sampled previously (where 2005 rebound plasma sequences were particularly low-diversity; mean = 0.025). Overall, these observations suggest that, while participant 4’s *replication-competent* reservoir sequences largely integrated around the time of ART initiation (as evidenced by the similar root-to-tip divergence measurements of the pre-ART sequences circulating in 2000 to those that rebounded in plasma in both 2004 and 2005), their total proviral pool, which also features defective proviruses, contains a substantial proportion of sequences that are far more ancestral than this (as evidenced by the abundance of proviral sequences deep in the tree, close to the root).

**Fig 9.**
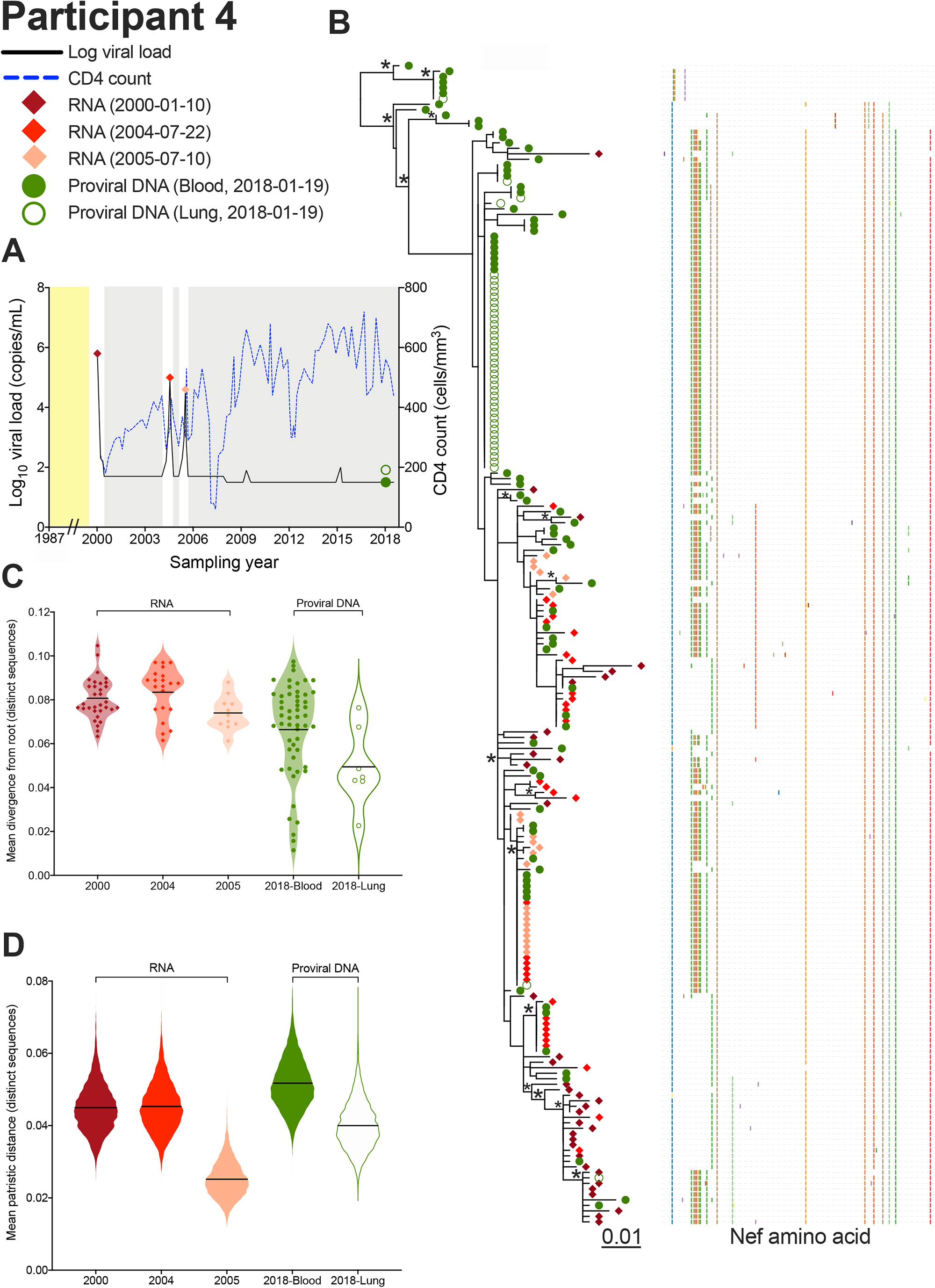
Interpreting proviral diversity in context of prior within-host HIV evolution: participant 4. (A) Plasma viral load (solid black line), CD4^+^ T-cell count (dashed blue line) and samples analyzed (colored diamonds and circles). Yellow shading denotes period following infection where no clinical information is available. Grey shading denotes periods of suppressive ART. (B) Highest likelihood phylogeny derived from Bayesian inference, rooted on the HIV reference strain HXB2. Scale shows estimated substitutions per nucleotide site. The adjacent plot shows amino acid diversity, with coloured ticks denoting non-synonymous substitutions with respect to the reference sequence at the top of the tree. Asterisks identify nodes supported by posterior probabilities ≥70%. (C) Mean divergence from the root of each distinct sequence, averaged over all 7,500 trees, stratified by sampling date and sample type. The black line represents the grand mean. (D) Within-host HIV sequence diversity, calculated as the mean patristic distance between all distinct sequences collected at that time point for that sample, where the values for each of the 7,500 trees are represented as a violin plot. The black line represents the grand mean.

## Discussion

The availability of matched blood and lung specimens from ART-suppressed participants undergoing bronchoscopy allowed us to investigate proviral diversity and compartmentalization across these sites. We recovered subgenomic proviral sequences from the lungs of all participants, confirming this organ as a site of long-term HIV persistence during ART. Proviral genetic composition however differed markedly between participants. First, identical sequence abundance (suggestive of clonally-expanded cell populations) ranged widely. Whereas a single proviral sequence dominated in the lung in some participants (e.g. 4 and 8), in others (e.g. 5) a specific sequence dominated in blood. In yet others (e.g. 1 and 7), few or no identical sequences were recovered. While this supports clonal expansion of latently-infected cells as a common driver of HIV persistence during ART in many individuals [14, 15, 51], it also reminds us that clonally-expanded populations are not present in everyone at all times. Identical sequences were not consistently enriched in one site compared to the other, and lung proviruses were not inherently more homogenous than blood, indicating that we cannot make generalizations regarding where clonally-expanded latently-infected cells are more likely to arise or persist in the body. In fact, overall within-host diversity of distinct blood and lung proviruses correlated significantly, indicating that proviral diversity in tissues generally reflects that in blood.

We also investigated the extent and type of blood/lung genetic compartmentalization, and whether it tends to present as distinct viral lineages across sites, or simply as differential proportions of clonally-expanded populations. We did this by analyzing distinct sequences *per compartment* (to detect the former) and including all sequences (to detect the latter). Importantly, no participant exhibited genetic compartmentalization so obvious that it was visually apparent in their phylogeny. When formal tests were applied however, two of nine individuals (22%, participants 2 and 6) showed significant evidence of distinct viral lineages across sites, though the extent of compartmentalization was modest (Table 2). When including all sequences, this number increased to four participants (44%) (Fig 7A). Of note, compartmentalization support for participant 6’s dataset increased substantially when all sequences were considered, indicating that their dataset featured both distinct lineages and differential distribution of identical sequences across sites. Overall, our results indicate that compartmentalization is generally limited, of modest magnitude, and when present it is often due to differential distributions of clonally-expanded populations. This observation is consistent with prior studies of viral diversity in lung [39, 40], female genital tract [34], testes [33], and lymph nodes [52].

The availability of archived plasma from two participants (4 and 6) allowed us to study on-ART proviruses in the context of pre-therapy viral diversity, revealing markedly different proviral dynamics. In participant 4, proviral sequences were more diverse and on average less divergent from the root than plasma sequences sampled just prior to ART initiation. Proviruses sampled on ART made up essentially all clades closest to the root, which was quite deep in the tree, suggesting that we had reconstructed back to a within-host MRCA that had existed quite some years ago. This in turn means that a substantial number of proviral sequences were more than 20 years old at the time of sampling in 2018. In contrast, participant 6’s tree root was not too much shallower than the least divergent sequences that circulated in plasma just before ART, suggesting that the reconstructed MRCA was relatively recent. Moreover, proviral sequences did not predominate near the root, and in fact were on average less diverse than the HIV RNA sequences that circulated in plasma just prior to ART initiation. Instead, this individual’s proviral pool more closely resembled the plasma HIV RNA sequences that rebounded and replicated during a 3-year treatment interruption between 2006-2009, where initial rebounding variants seemed to be on average somewhat older than the ones that circulated just prior to ART.

These findings are notable in light of recent observations that proviruses persisting on ART are typically enriched in sequences that integrated around ART initiation [10, 12, 49], though some individuals preserve larger numbers of ancestral sequences [7, 8, 49, 50]. Differential pre-ART proviral dynamics explain some of these differences: while HIV sequences continually enter the reservoir during untreated infection, proviral turnover during this time is relatively rapid compared to during ART, which slows proviral turnover, thereby creating an environment more favorable to proviral persistence [49, 53–55]. Indeed, recent studies have estimated the average half-life of persisting proviruses to be on the order of only ∼1 year during untreated infection [11, 12, 53], which means that most proviruses that enter the reservoir during this time will be “cleared” from the body by the time ART is initiated. Pre-ART proviral turnover is slower in some individuals however [49, 53], allowing a greater number of ancestral sequences to persist long-term. Participant 4, on-ART proviral pool contained a substantial portion of ancestral proviruses, falls into this latter category. Participant 6, by contrast, only exhibited one provirus that appeared more ancestral than any HIV RNA sequence circulating just prior to ART, and only marginally so. We hypothesize that this is due to a combination of two factors: first, a more typically rapid turnover rate during untreated infection that facilitated the elimination of ancestral sequences, and secondly the substantial re-seeding of their reservoir during the 2006-2009 treatment interruption. We acknowledge that these interpretations depend in part on tree root placement, which is inherently uncertain (especially when outgroup rooting [56, 57]). Nevertheless, the observation that participant 4’s blood proviruses are more diverse than the pre-ART plasma sequences, while the reverse is true for participant 6, is independent of tree root placement.

Our study has some additional limitations. Because we performed subgenomic sequencing, we cannot definitively classify identical sequences as being from clonally-expanded cells (full-genome sequencing plus integration site determination would be required to do this). Also, many sequences are likely not from intact proviruses since most proviruses persisting during ART harbor large deletions [58–60]. Intact subgenomic sequences, however, are still appropriate for inferring evolutionary relationships and testing for compartmentalization. This is because large deletions arise, likely in a single step, during reverse transcription of HIV RNA genomes that have newly infected a cell (this is supported by the observation that deletions reproducibly occur at locations flanked by identical sequence motifs [59–61]). As such, proviral regions that are unaffected by the deletion (or by other processes such as hypermutation) still reflect the sequence of the original infecting virus and are therefore appropriate for evolutionary inference. Indeed, the majority of within-host proviral diversity and compartmentalization studies are performed on subgenomic sequences (e.g. [3, 7, 9, 33, 50]. As biological material was limited, we isolated proviruses directly from blood and BAL cell pellets, so the cell types that harbored these sequences remain unknown. Our isolation of identical sequences in lung (suggestive of clonal expansion) suggests that these are derived from CD4^+^ T-cells (as macrophages are non-dividing [62]). For four participants (3, 5, 7 and 9) we isolated fewer than 10 intact proviral sequences from the lung, which limited our ability to detect compartmentalization. This suggests that lung proviral load was low in these individuals, though we had insufficient material to confirm this. Nevertheless, our recovery of diverse lung proviruses from all participants, which reflected the overall blood proviral diversity, indicates that blood proviral diversity mirrors that in tissues.

Our study has some strengths and novel aspects. Our use of Bayesian approaches [63] to generate a distribution of within-host phylogenies per participant allowed us to account for the inherent uncertainty in phylogenetic inference. Application of multiple compartmentalization tests allowed us to classify datasets as compartmentalized (or not) with more certainty. The availability of archived plasma from as long as 18 years prior to proviral sampling allowed us to interpret proviral diversity in context of HIV’s within-host evolutionary history for two participants, revealing marked differences in proviral longevity.

In summary, our results reveal that lung proviral diversity in individuals receiving long-term suppressive ART broadly reflects that in blood. Genetic compartmentalization across these two sites was not the norm, and when present it was of modest magnitude, and just as likely to be driven by differential clonal expansion across sites rather than the presence of distinct lineages. More broadly, our study highlights the genetic complexity of HIV proviruses persisting in lung and blood during long-term ART, and above all the uniqueness of each individual’s proviral composition and dynamics. Individualized strategies for HIV remission and cure strategies may be needed to overcome these challenges.

## Materials and methods

### Ethics statement

This study was approved by the Providence Health Care/University of British Columbia and Simon Fraser University research ethics boards. All participants provided written informed consent.

### Study participants and sampling

We studied paired buffy coat (“blood”) and bronchoalveolar lavage (BAL) (“lung”) specimens from nine participants with HIV, eight males and one female, who had undergone bronchoscopy at St. Paul’s Hospital in Vancouver, Canada. Participants were a median 60 (IQR, 51-62) years of age, and had maintained viremia suppression on ART for a median of 9 years (IQR, 4-13) at the time of sampling. Two participants (8 and 9) had been diagnosed with chronic obstructive pulmonary disease, the remainder had no documented overt respiratory symptoms or lung disease. Buffy coats had been separated from whole blood as described previously [64], and an estimated 5-10 million peripheral blood mononuclear cells (PBMC) were studied presently. BAL was performed according to previously described methods [65], spun down to collect cell pellets and resuspended in 1mL Cytolyt solution (Cytyc, Marlborough, MA); the full 1mL was studied presently. Buffy coat and BAL aliquots were stored at -80°C until further use. Longitudinal pre-ART plasma samples were also available from two participants (4 and 6).

### HIV amplification and sequencing

DNA was extracted from buffy coat and BAL using the QIAamp DNA Mini Kit (Qiagen). Single-genome amplification of a subgenomic HIV region (*nef*) was performed using primers designed to amplify major HIV subtypes [33, 50]. Briefly, genomic DNA extracts were endpoint diluted such that <30% of the resulting nested polymerase chain reactions (PCR), performed using an Expand High Fidelity PCR system (Roche), would yield an amplicon. First round PCR primers were Nef8683F_pan (forward; TAGCAGTAGCTGRGKGRACAGATAG) and Nef9536R_pan (reverse; TACAGGCAAAAAGCAGCTGCTTATATGYAG). Second round PCR primers were Nef8746F_pan (forward; TCCACATACCTASAAGAATMAGACARG) and Nef9474R_pan (reverse; CAGGCCACRCCTCCCTGGAAASKCCC). Negative controls were included in every run. For the participants with archived plasma available, nucleic acids were extracted using the BioMerieux NucliSENS EasyMag system (BioMerieux, Marcy-l’Étoile, France). Next, cDNA was generated using NxtScript reverse transcriptase (Roche) using the reverse primer Nef9536R_pan, after which the cDNA was endpoint diluted and single-genome-amplified as described above. Amplicons were sequenced on a 3730xl automated DNA sequencer using BigDye (v3.1) chemistry (Applied Biosystems). Chromatograms were base called using Sequencher (v5.0/v5.4.6) (GeneCodes).

### Sequence alignment and phylogenetic inference

Sequences exhibiting genetic defects (including large insertions/deletions), nucleotide mixtures, hypermutation (identified using Hypermut [66]) were excluded, as were sequences exhibiting evidence of within-host recombination (identified using RDP4 v4.1) [67]. Sequences were aligned in a codon-aware manner using MAFFT v7.471 [68]. Alignments were inspected and manually edited in AliView v1.26 [69]. A maximum-likelihood between-host phylogeny was inferred using IQ-TREE2 [70] following automated model selection using ModelFinder [71]. Branch support values were derived from 1,000 bootstraps.

Within-host phylogenies were inferred using Bayesian approaches as follows. First, each within-host nucleotide alignment was reduced to distinct sequences only (where an identical sequence was found in both lung and blood, only one instance was retained). The best-fitting substitution model for each dataset was determined using jModelTest v2.1.10 [72]. Next, Markov chain Monte Carlo (MCMC) methods were used to build a random sample of phylogenies per participant. Two parallel runs with MCMC chains of five million generations, sampled every 1,000 generations, were performed in MrBayes, v3.2.7 [73] using the best-fitting substitution model and model-specific or default priors. Convergence was assessed by ensuring that the deviation of split frequencies was <0.03, that the effective sampling size of all parameters was ≥200, and through visual inspection of parameter traces in Tracer, v1.7.2 [74]. The first 25% of iterations were discarded as burn-in, yielding 7,500 phylogenies per participant.

Identical sequences were then grafted back on to these trees using the add.tips function in R package phangorn, v2.8.1 [75] to generate two sets of trees: first, trees containing distinct sequences *per compartment* (here, any sequence that was observed in both blood and lung was added back to the tree if it had been initially removed from that compartment) and second, trees containing all within-host sequences collected. Phylogenies were plotted using the R (v 4.1.2) package ggtree, version 3.21. The tree with the highest likelihood, midpoint rooted in FigTree (v1.4.4) (http://tree.bio.ed.ac.uk/software/figtree/) is displayed for each participant. Node support values are derived from Bayesian posterior probabilities from the consensus tree.

For participants 4 and 6, for whom pre-ART plasma HIV RNA sequences were available, MCMC runs were additionally performed on alignments of proviral (blood/lung) and plasma HIV RNA sequences as described above, and trees were rooted as described in the results. For participant 6, the root location that maximized the correlation between root-to-tip distances and sampling time of plasma sequences collected in 2006 and 2009 was identified using the in-house software phylodating (https://bblab-hivresearchtools.ca/django/tools/phylodating/ and [7]). For participant 4, outgroup rooting was performed using the evolutionary placement algorithm in RaxML, implemented with a custom script in R [76].

### Genetic compartmentalization and statistical analyses

Compartmentalization was assessed using two genetic distance-based tests: Analysis of Molecular Variance (AMOVA) [45] and Hudson, Boos and Kaplan’s nonparametric test for population structure (KST) [46], and two tree-based tests: the Slatkin-Maddison test (SM) [47] and the Correlation Coefficients (CC) test [48]. KST was run in Hyphy, v2.5.2 [77] using the TN93 genetic distance matrix. AMOVA was implemented in the R package pegas, v1.1 [78]. For these tests, statistical significance was assessed via 1,000 permutation tests. The tree-based tests were applied to all 7,500 phylogenies inferred per participant, and the output statistics were averaged across all trees. This was done using the R package slatkin.maddison, v0.1.0 for the SM test [79], and using a custom R script for the CC test. A within-host dataset was classified as compartmentalized when at least two tests returned statistically significant results. Tests were first performed on distinct sequences per compartment, and again on all within-host sequences.

Blood and lung proviral diversity was assessed using two methods. For each phylogeny, we first computed the mean within-host patristic distance between all pairs of distinct sequences per compartment, and then calculated the mean over all 7,500 trees (i.e., *grand mean patristic distances*). Second, we summed the edge lengths of all distinct sequences per compartment in each tree, and then calculated the mean over all 7,500 trees (i.e., *mean phylogenetic diversity*) [80]. Both metrics were computed using custom R scripts. All other statistical analyses were performed in Prism, v9.0 (GraphPad Software). All compartmentalization tests of significance were one tailed (all others two tailed) with p < 0.05 denoting statistical significance unless otherwise indicated.

### Data deposition

The nucleotide sequences reported in this paper are available in GenBank (accession numbers: proviral DNA: OM963156 – OM964037; HIV RNA: OM964038 – OM964339).

## Acknowledgements

We sincerely thank the study participants without whom this research would not be possible. We thank Mark Brockman for helpful discussions. We thank Hanwei Sudderuddin, Hope Lapointe and Sarah Speckmaier for technical assistance.

This work was supported by the Canadian Institutes of Health Research (CIHR) through a project grant (PJT-159625 to Z.L.B. and J.B.J.) and a focused team grant (HB1-164063 to Z.L.B.). This work was supported by the Martin Delaney ‘BELIEVE’ Collaboratory (NIH grant 1UM1AI26617 to Z.L.B.), which is supported by the following NIH Co-Funding and Participating Institutes and Centers: NIAID, NCI, NICHD, NHLBI, NIDA, NIMH, NIA, FIC, and OAR. Research reported in this publication was also supported by the National Institute of Allergy and Infectious Diseases of the National Institutes of Health under Award Number UM1AI164565 (Martin Delaney ‘REACH’ collaboratory; to Z.L.B.). The content is solely the responsibility of the authors and does not necessarily represent the official views of the National Institutes of Health. A.S. and B.R.J. are supported by CIHR Doctoral Research Awards. J.M.L is supported by a Health Professional Investigator Award from the Michael Smith Foundation for Health Research and by a Tier 2 Canada Research Chair in Translational Airway Biology. Z.L.B. is supported by a Scholar Award from the Michael Smith Foundation for Health Research.

